# Order of Message and Address Domain Engagement Determines Productive β-Endorphin Binding to the μ-Opioid Receptor

**DOI:** 10.64898/2026.02.06.704327

**Authors:** Maciej P. Ciemny, Sebastian Kmiecik

## Abstract

Understanding *β*-endorphin binding to the μ-opioid receptor (*μ*OR) is crucial for designing safer analgesics. The peptide comprises a message domain mediating activation and an address domain conferring selectivity. Using 1,000 independent CABS-dock simulations, without prior binding-site knowledge, we analysed binding trajectories to compare alternative binding pathways. Message-first binding is most frequently sampled but rarely reaches native-like structures (5.0%). In contrast, address-first binding occurs less often yet shows a 3.8-fold higher success rate (18.8%, p < 0.001). These results refine the message–address model and suggest that early address-domain engagement promotes productive μOR binding.

## INTRODUCTION

Opioid receptors (ORs) are critical targets for analgesic development, yet current drugs cause severe side effects including respiratory depression and addiction.^1^ Among endogenous opioid peptides, *β*-endorphin exhibits exceptional μ-opioid receptor (*μ*OR) affinity through a message-address architecture:^2^ the N-terminal YGGF motif (“message”, residues 1-4) drives receptor activation, while the C-terminal sequence (“address”, residues 11-21) provides selectivity. Recent cryogenic electron microscopy structure (PDB: 8F7Q)^1^ resolved residues 1-21 bound to *μ*OR-inhibitory G-protein complex, providing unprecedented molecular detail of this interaction.

Understanding binding may inform biased ligand design, as different domains could preferentially activate G-protein versus *β*-arrestin pathways.^3^ However, computational docking of flexible peptides to G-protein coupled receptors (GPCRs) remains challenging due to high conformational flexibility and membrane environment effects.^4^ While methodological advances enable peptide-GPCR modeling,^5–7^ systematic validation has focused on shorter peptides lacking explicit membrane engagement.^4^

What remains unclear is whether the canonical assumption that address-domain recognition initiates productive binding holds for long, flexible peptides such as β-endorphin, or whether alternative orders of domain engagement may more effectively stabilize native-like receptor complexes. Here we investigate *β*-endorphin-*μ*OR binding using CABS-dock^8^ with membrane modeling^9^ through 1,000 independent simulations, enabling a comparative analysis of distinct binding pathways and their structural outcomes.

## MATERIALS AND METHODS

### Structure Preparation

The *μ*OR-*β*-endorphin complex structure was extracted from 8F7Q PDB^10^ entry. All components aside from the receptor and the peptide (inhibitory G protein, nanobody, cholesterol hemisuccinate, waters, ions) were removed.

### Molecular Docking

CABS-dock^5,8,11^ with membrane extension^9^ was used to perform flexible docking using coarse-grained representation and replica-exchange Monte Carlo (MC) sampling (10 replicas per simulation). CABS-dock is a well-established global docking approach that does not require prior knowledge of the binding site and predicts peptide structure de novo, starting from random conformations and random positions relative to the receptor.^5,8,11^ In addition to linear peptides, the CABS-dock framework has recently been successfully applied to the flexible docking of cyclic peptides, demonstrating its applicability to structurally constrained and challenging peptide systems.^12^

Beyond structure prediction, CABS-dock has been successfully applied to investigate protein–peptide binding mechanisms, peptide folding upon binding, and interactions involving intrinsically disordered fragments.^13,14^ The applicability of the CABS force field to model protein and peptide dynamics has been extensively reviewed elsewhere. ^15,16^

The membrane was modeled as a continuous hydrophobic region by introducing additional terms in the scoring function.^9^ We conducted 1,000 independent simulations without binding site knowledge nor peptide bound structure, generating 10,000 trajectories (1,000 frames each, ∼10 million snapshots total).

### Classification Criteria

An interaction complex was defined with thresholds for each of the peptide domains: Message root-mean squared deviation, RMSD <10 Å, Address RMSD<15 Å (which mark 10th percentile distributions for these domains); Message contacts 37.5%, Address contacts 20.0% (75th percentile within binding-relevant frames, RMSD < 10/15 Å). RMSD-contact recovery correlations validated RMSD as a reliable binding quality proxy across all domains (Message r = −0.707, Address r = −0.535, Total r = −0.686; all p < 0.001). RMSD was used as its continuous character allows for more detailed analysis than contacts which are discrete by definition. Interaction complexes were identified at MC replica level by sustained 20-frame windows meeting both RMSD and contact criteria within a single replica trajectory. Native-like binding was defined at the MC replica level as mean total RMSD < 5 Å over 100 frames within replica trajectory (mirroring the standardized Critical Assessment of Prediction of Interactions “acceptable” quality threshold).^17^ Frame indices in replica trajectories reflect Monte Carlo sampling progression and are not interpreted as physical time.

Trajectories were classified as: message-first (Δt > 50 frames), address-first (Δt < −50 frames), Message-Only, Address-Only, Simultaneous (|Δt| ≤ 50 frames), or Unbound, where Δt = Address binding frame − Message binding frame. For mechanistic analysis, Simultaneous events were grouped with message-first based on: (i) 89% showed initial Message contact within ±10 frames; (ii) success rates matched message-first (7.3% vs 4.5%, p = 0.18) not address-first (18.8%, p < 0.001); (iii) similar contact pattern dynamics. All analyses were additionally performed with Simultaneous trajectories treated as a separate class, yielding no change in the relative ranking of pathway productivity (data not shown).

Contact maps calculated using CABS side chain pseudoatoms positioned at geometric centers of side chain heavy atoms.^18^ Side chain orientations determined by local backbone geometry. The residue-residue contacts were defined as pseudoatom distance < 6.5 Å.

## RESULTS AND DISCUSSION

### Dataset Overview and Pathway Heterogeneity

We analysed 10 000 replica trajectories generated from 1 000 independent CABS-dock simulations using replica-exchange Monte Carlo sampling. Of these, 2 334 replica trajectories formed interaction complexes that met the sustained binding criteria. Despite clear heterogeneity in binding pathways, the most frequently sampled pathway did not correspond to the most productive one. Initial binding events were dominated by the message-first pathway (n = 1 485, 63.6%), followed by the address-first pathway (n = 499, 21.4%), with the remaining trajectories remaining unbound. Thus, the two productive pathways coexist in an approximate 3.0:1 ratio (Figure 1A). The prevalence of message-first binding is consistent with the smaller conformational search space of the message domain (4 residues versus 11 residues in the address domain). The structural context of the message–address pharmacophore used throughout the analysis is shown in Figure 1C.

**Figure 1.**
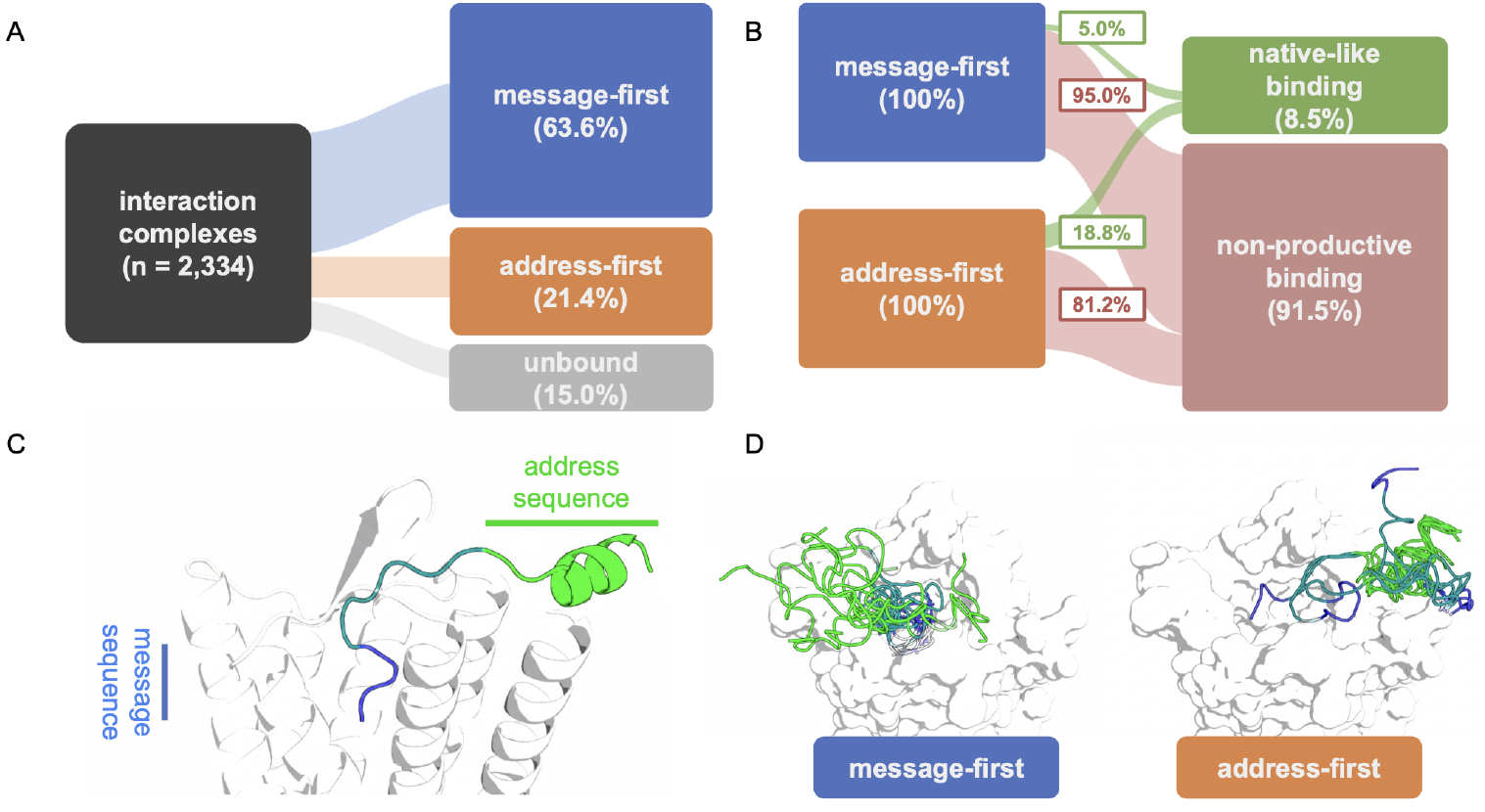
Distinct binding pathways of β-endorphin to the μ-opioid receptor. (A) Distribution of interaction complexes among binding pathways across replica trajectories, showing the relative prevalence of message-first, address-first, and unbound events. (B) Conditional success rates of native-like binding within each pathway, highlighting the inverse relationship between pathway prevalence and productive binding. Flow widths are normalized independently for message-first and address-first trajectories. (C) Reference structure of the μ-opioid receptor bound to β-endorphin (PDB: 8F7Q), illustrating the message–linker–address domain organization used throughout the analysis. (D) Representative intermediate structures sampled along message-first (left) and address-first (right) trajectories that subsequently reach native-like binding.

### Pathway Prevalence versus Productive Binding

Although message-first pathways are sampled most frequently, they exhibit markedly lower productivity. Address-first replica trajectories reached native-like binding in 94 of 499 cases (18.8%, 95% CI: 15.6–22.5%), whereas message-first trajectories succeeded in only 74 of 1,485 cases (5.0%, 95% CI: 4.0–6.2%). This corresponds to a 3.8-fold higher conditional success rate for address-first pathways (z = 9.62, p < 0.001; Figure 1B). Thus, the pathway most frequently sampled in the simulations is not the most likely to yield native-like complexes. The results indicate that early engagement of the address domain is associated with a higher proportion of sampled conformations that resemble the native complex, whereas message-first recognition more frequently populates non-native binding arrangements. Throughout this study, ‘productive binding’ refers exclusively to structural convergence toward the cryo-EM reference complex and does not imply receptor activation or signaling efficacy. The analysis is limited to structural sampling within the Monte Carlo framework.

The two pathways exhibit symmetrically inverted profiles, in which the threefold prevalence advantage of message-first binding is offset by the 3.8-fold higher success rate of address-first binding, resulting in comparable overall contributions to the native-like structure population. To assess whether these productive trajectories correspond to structurally accurate binding modes, we next evaluated the structural quality and contact patterns of the sampled complexes. The relative differences between message-first and address-first pathways remain unchanged across reasonable variations of RMSD and contact thresholds (Figure S3).

### Structural Quality and Contact Analysis

The computational ensemble generated by the CABS-dock protocol exhibits extensive sampling of the receptor surface while consistently capturing near-native binding configurations (Figure S1). Minimum RMSD values reached 0.94 Å for the message domain, 2.29 Å for the address domain, and 2.58 Å for the full peptide, with native contact recovery reaching complete recovery for individual domains and 77.8% for the full peptide–receptor interface. These metrics demonstrate that the simulations reliably sample structurally accurate binding modes, providing a robust foundation for pathway-level comparisons.

To identify molecular determinants underlying productive binding, we analysed peptide–receptor contact frequencies across all frames from replica trajectories that achieved native-like complexes (n = 168). This analysis recapitulated 10 of 11 experimentally reported interaction hotspots at receptor positions 126, 132, 149, 150, 153, 218, 312, 320, 324, and 328 (Figure 2). The remaining hotspot at residue 135 exhibited lower-than-expected contact frequency, likely reflecting limitations of the coarse-grained force field in describing this critical orthosteric interaction.

**Figure 2.**
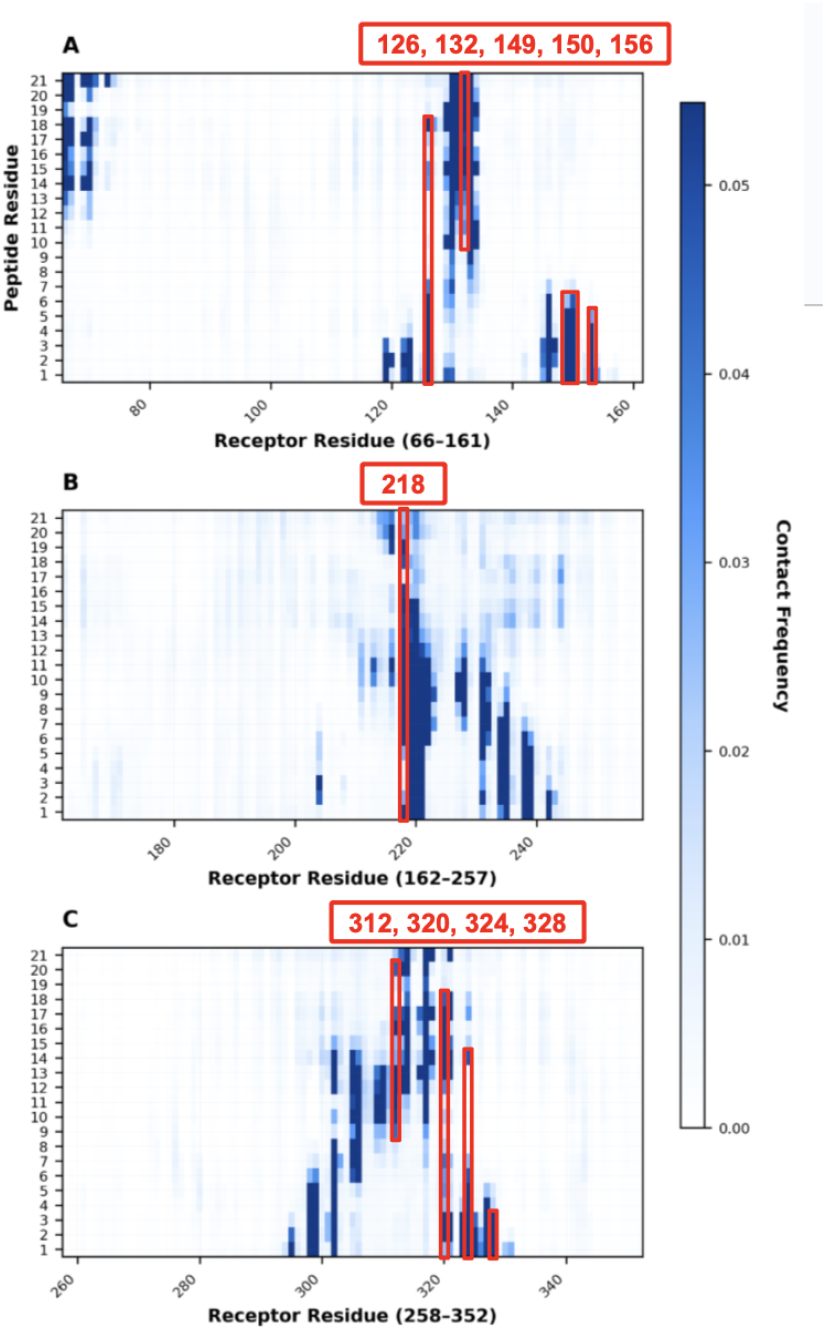
Peptide–receptor contact frequency map from productive docking trajectories. Contact frequencies were calculated across all frames from replica trajectories that achieved native-like binding (mean total RMSD < 5 Å over 100 frames), excluding non-productive interaction complexes. Contacts were defined as CABS side chain center of mass distances < 6.5 Å. Red boxes indicate experimentally validated native contact hotspots that are recapitulated by the simulations, shown only at positions with significant contact frequency. The receptor is segmented into three panels (A–C) for visualization clarity.

## CONCLUSIONS

Large-scale docking simulations reveal that β-endorphin binding to the μ-opioid receptor proceeds through two distinct pathways that differ markedly in their likelihood of forming native-like complexes. The message-first pathway is most frequently sampled during early stages of structural sampling but often leads to non-native binding arrangements, with only 5.0% of trajectories reaching native-like states. In contrast, address-first binding occurs less frequently yet shows a substantially higher probability of yielding native-like complexes (18.8%). Together, these results refine the classical message–address paradigm by demonstrating that the order of domain engagement, rather than the prevalence of initial message-domain recognition, is a key determinant of productive μOR binding.

Beyond mechanistic insight, this pathway differentiation has direct implications for peptide design. Strategies focused exclusively on optimizing message-domain recognition may preferentially enrich frequently sampled but structurally unstable binders. In contrast, promoting early and sustained engagement of the address domain—through conformational preorganization, linker optimization, or screening criteria that prioritize binding stability over initial encounter prevalence - may improve the likelihood of productive receptor activation. Structural analysis of productive trajectories supports this view by recapitulating experimentally validated receptor contact hotspots and near-native binding configurations.

These conclusions should be interpreted within the limits of the coarse-grained representation, which enables extensive sampling at the expense of atomic detail, as well as the approximate treatment of receptor flexibility and membrane effects. Future studies incorporating explicit receptor dynamics and membrane models will be important for quantifying how lipid interactions and conformational constraints further modulate address-domain engagement and pathway selection. Within these bounds, the present work provides a framework for linking binding pathway order, structural stabilization, and pharmacophore architecture in peptide–GPCR recognition.

## ABBREVIATIONS

OR: opioid receptor
GPCR: G-protein coupled receptor
PDB: protein data bank
RMSD: root-mean-squared deviation
μOR: μ-opioid receptor

## DATA AVAILABILITY

The cryo-EM structure of the *μ*-opioid receptor in complex with *β*-endorphin (residues 1–21) used as the reference for this study is deposited in the Protein Data Bank under accession code 8F7Q. The CABS-dock standalone package is available at https://bitbucket.org/lcbio/cabsdock/; the membrane-enabled extension used in this work can be accessed at https://github.com/mcmny/cabs-dock-membrane. Representative output trajectories from 1,000 independent simulations, representative structures used for visualization in Figure 1D, data analysis scripts, and processed data files underlying Figures 1A, 1B, 2, S1, and S2 are deposited at https://doi.org/10.5281/zenodo.18495710.

## ASSOCIATED CONTENT

### Supporting Information

Additional methodological details and figures: Figure S1 (structural quality metric distributions across full ensemble), Figure S2 (evolution of contacts during message domain binding).

## AUTHOR INFORMATION

### Corresponding Author

Sebastian Kmiecik - Biological and Chemical Research Center, Faculty of Chemistry, University of Warsaw, Pasteura 1, 02-093 Warsaw, Poland.

### Author Contributions

The manuscript was written through contributions of all authors. All authors have given approval to the final version of the manuscript.

### Notes

Any additional relevant notes should be placed here.

## ACKNOWLEDGMENT

Maciej P. Ciemny has been supported by the National Science Centre, Poland (Preludium 2018/29/N/NZ1/01196).

