## Supplementary Information for "Order of Message and Address Domain Engagement Determines Productive β-Endorphin Binding to the μ-Opioid Receptor"

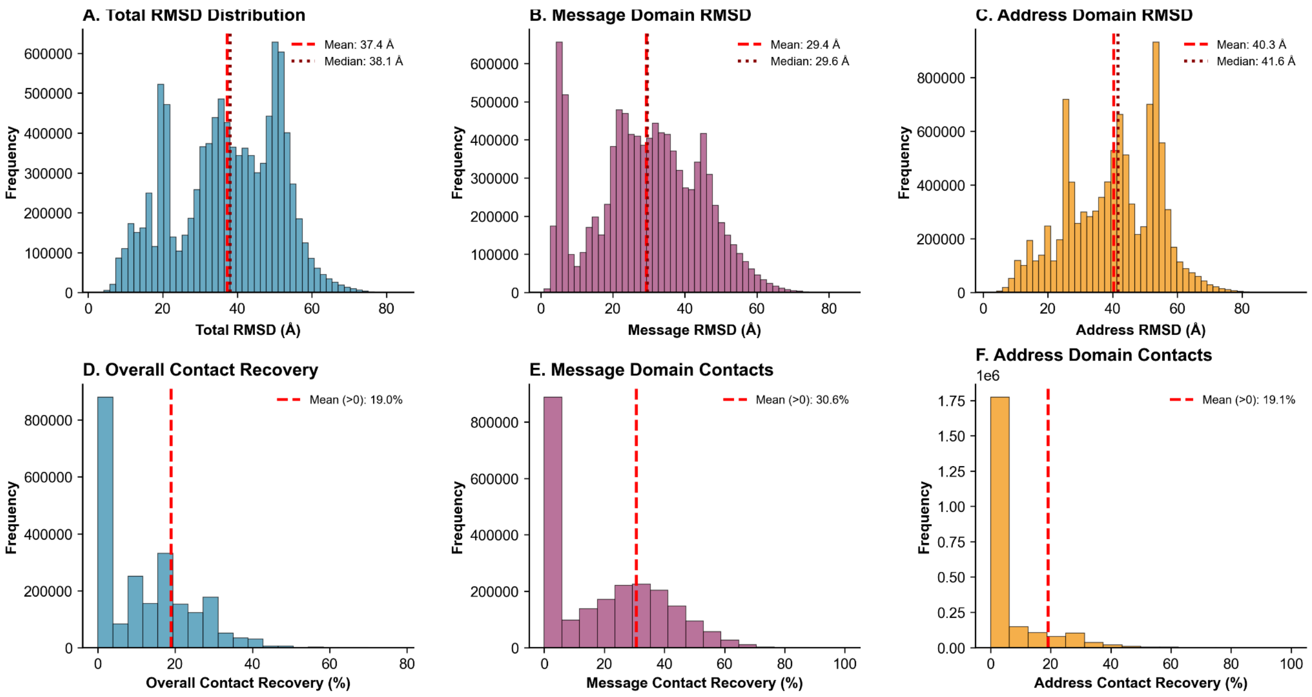


**Figure S1. Distribution of Structural Quality Metrics Across the Entire Simulation Ensemble.** Histograms showing RMSD and contact recovery distributions across all frames from the complete computational ensemble (10,000 replica trajectories × 1,000 frames = ~10 million total frames). RMSD distributions for (A) total peptide, (B) message domain (residues 1-4), and (C) address domain (residues 11-21), respectively. Red dashed lines indicate mean values; dotted lines show medians. The high mean values (Total: 37.4 Å, Message: 29.4 Å, Address: 40.3 Å) reflect the blind, unbiased nature of the global search across the entire receptor surface, with individual frames sampling the full conformational space without binding site constraints. Native contact recovery distributions for the (D) entire peptide-receptor interface, (E) message domain contacts, and (F) address domain contacts, respectively. Red dashed lines show the mean contact recovery calculated only for frames with non-zero contacts (>0). The broad distributions demonstrate extensive conformational sampling at the frame level across all 10,000 replica trajectories.

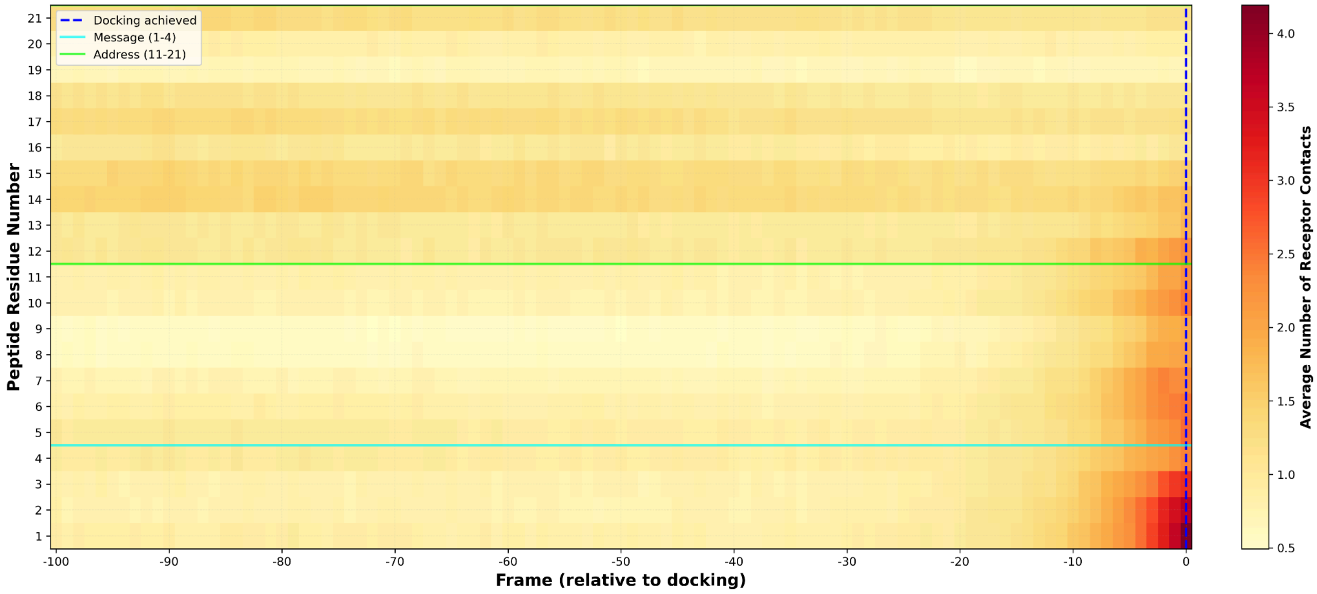


**Figure S2. Evolution of Peptide-Receptor Contacts Leading to Message Domain Docking.** Heatmap showing average receptor contact counts per peptide residue as a function of frame number relative to message domain docking achievement (Frame 0, defined as the first frame within a replica trajectory in which message RMSD < 5 Å is achieved). Data represents frame-level analysis averaged across all replica trajectories that achieved message docking (n = replicas meeting message RMSD < 5 Å criterion). Color intensity represents the mean number of receptor contacts per residue per frame (yellow to dark red scale). Horizontal lines delineate message domain (cyan, residues 1-4) and address domain (green, residues 11-21). The blue dashed vertical line marks Frame 0 (docking achievement frame). The temporal contact profile reveals minimal pre-docking organization at the frame level: frames preceding message docking (−100 to −10 frames before Frame 0) show uniformly low contact levels across all peptide residues, indicating limited productive engagement during the approach phase within individual replica trajectories. Contact formation occurs abruptly, with message residues 1-4 showing sharp increases (yellow to red transition) coincident with RMSD threshold crossing at Frame 0. Address domain residues (11-21) exhibit modest contact increases, with residues 12 and 21 showing elevated engagement, suggesting partial coordination with message binding. The lack of gradual contact accumulation in pre-docking frames indicates that successful message engagement occurs through "collision-and-capture" rather than progressive approach, with the transition happening over ≤10 frames within individual replicas.

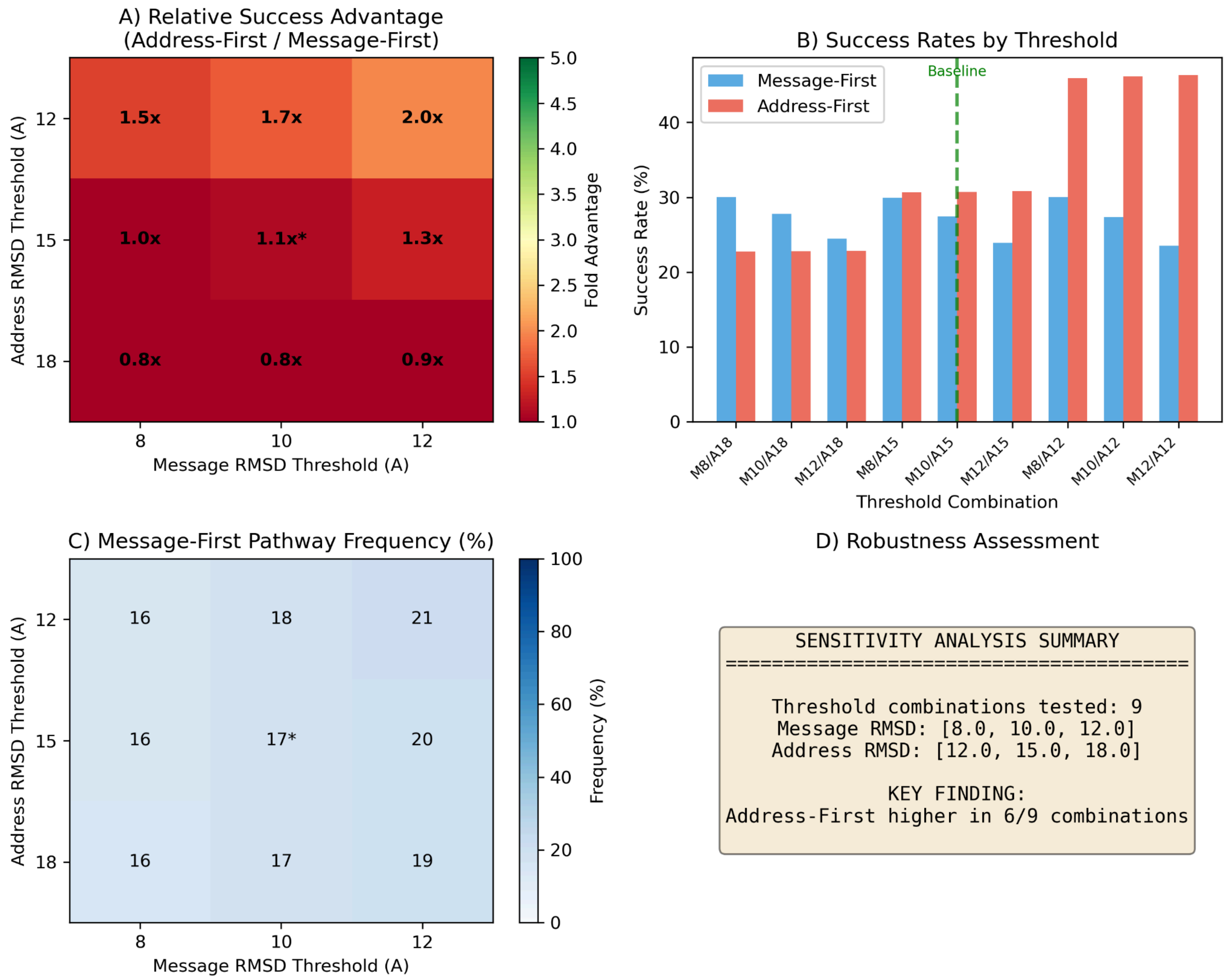


**Figure S3. Sensitivity analysis of pathway classification to RMSD threshold selection.** Sensitivity analysis across 9 threshold combinations revealed threshold-dependent pathway advantages. With physically meaningful Address thresholds (≤15 Å), Address-First pathways showed 1.0–2.0× higher success rates. The 18 Å Address threshold represents a control condition: at this distance, structures lack specific receptor contacts and represent non-specific association rather than binding. The loss of Address-First advantage at 18 Å (0.8–0.9×) confirms that the observed pathway differences reflect genuine binding rather than classification artifacts. The consistent Address-First advantage across all biologically meaningful thresholds (12–15 Å) demonstrates robustness of the core finding. (A) Heatmap of relative success advantage (Address-First / Message-First) across threshold combinations; asterisk indicates baseline thresholds used in main analysis. (B) Success rates for both pathways across all combinations; green dashed line indicates baseline. (C) Message-First pathway frequency across threshold combinations. (D) Summary of robustness assessment. MF = Message-First pathway (including Message-Only and Simultaneous); AF = Address-First pathway (including Address-Only).
